# Quantifying relative virulence: When μ_max_ fails and AUC alone just isn’t enough

**DOI:** 10.1101/2020.04.05.026013

**Authors:** Ruben Michael Ceballos, Carson Len Stacy

## Abstract

One of the more challenging aspects in quantitative virology is quantifying relative virulence between two (or more) viruses that have different replication dynamics in a given susceptible host. Host growth curve analysis is often used to detail virus-host interactions and to determine the impact of viral infection on a host. Quantifying relative virulence using canonical parameters such as maximum specific growth rate (μ_max_) can fail to provide accurate information regarding experimental infection, especially for non-lytic viruses. Although area-under-the-curve (AUC) can be more robust by through calculation of a percent inhibition (PI_AUC_), this metric can be sensitive to limit selection. In this study, using empirical and extrapolated data from Sulfolobus Spindle-shaped Virus (SSV) infections, we introduce a novel, simple metric that is proven to be more robust and less sensitive than traditional measures for determining relative virulence. This metric (*I*_*SC*_) more accurately aligns biological phenomena with quantified metrics from growth curve analysis to determine trends in relative virulence. It also addresses a major gap in virology by allowing comparisons between non-lytic single-virus/single-host (SVSH) infections and between non-lytic versus lytic virus infection on a given host. How *I*_*SC*_ may be applied to polymicrobial infection – both coinfection of a host culture and superinfection of a single cell with more than one virus (or other pathogen type) is a topic of ongoing investigation.

## Introduction

Quantifying relative virulence (*V*_*R*_) is challenging when comparing viruses that exhibit different replication dynamics in a given host. Although ***host growth curve analysis*** is often used to detail virus-host interactions and to determine the level of detriment a virus levies on host growth, assessing *V*_*R*_ via canonical measures of fitness, such as maximum specific growth rate (*μ*_*max*_) [1], can fail to accurately describe experimental infection datasets [2], especially for non-lytic viruses. In non-lytic virus systems, progeny virions are released via *budding* rather than gross cell lysis and growth curves for hosts infected with non-lytic viruses can exhibit non-canonical growth profiles, including absence of a *lag* phase and very brief *exponential* growth followed by a prolonged period of non-exponential (but positive) growth prior to *stationary* phase.

Using empirical and extrapolated data from Sulfolobus Spindle-shaped Virus (SSV) infections, we introduce a novel, simple metric that overcomes limitations of traditional growth curve analysis when quantifying relative virulence between two viruses independently infecting a common host. This approach (*viz:* Stacy-Ceballos equations; *see* Eqs. 3, 4) more accurately aligns biological phenomena with quantified metrics for *V*_*R*_ and addresses a major gap in virology by allowing comparisons between non-lytic and lytic virus infections.

For this report, *maximum specific growth rate* (*μ*_*max*_) and percent inhibition based on AUC (PI_AUC_) calculations were used to determine *V*_*R*_ for strains of Sulfolobus Spindle-shaped Virus (SSV). SSVs are non-lytic double-stranded DNA viruses that infect species of the family *Sulfolobaceae* – a group of hyperthermophilic archaea. *V*_*R*_ across three SSVs was assessed by comparing parameters between growth curves from host cultures, each of which was infected with one of three viruses: SSV1 [3], SSV2 [4], and SSV8 [5] – in single-virus/single-host infections on *Sulfolobus* strain Gθ [6,7]. Absorbance data (a proxy for cell density) were fit with Logistic and Gompertz models. Both model types exhibit similar *goodness of fit* (Fig. 1A, B); however, Gompertz [8] is preferred for analyzing diseased cells [9,10].

**Fig 1.**
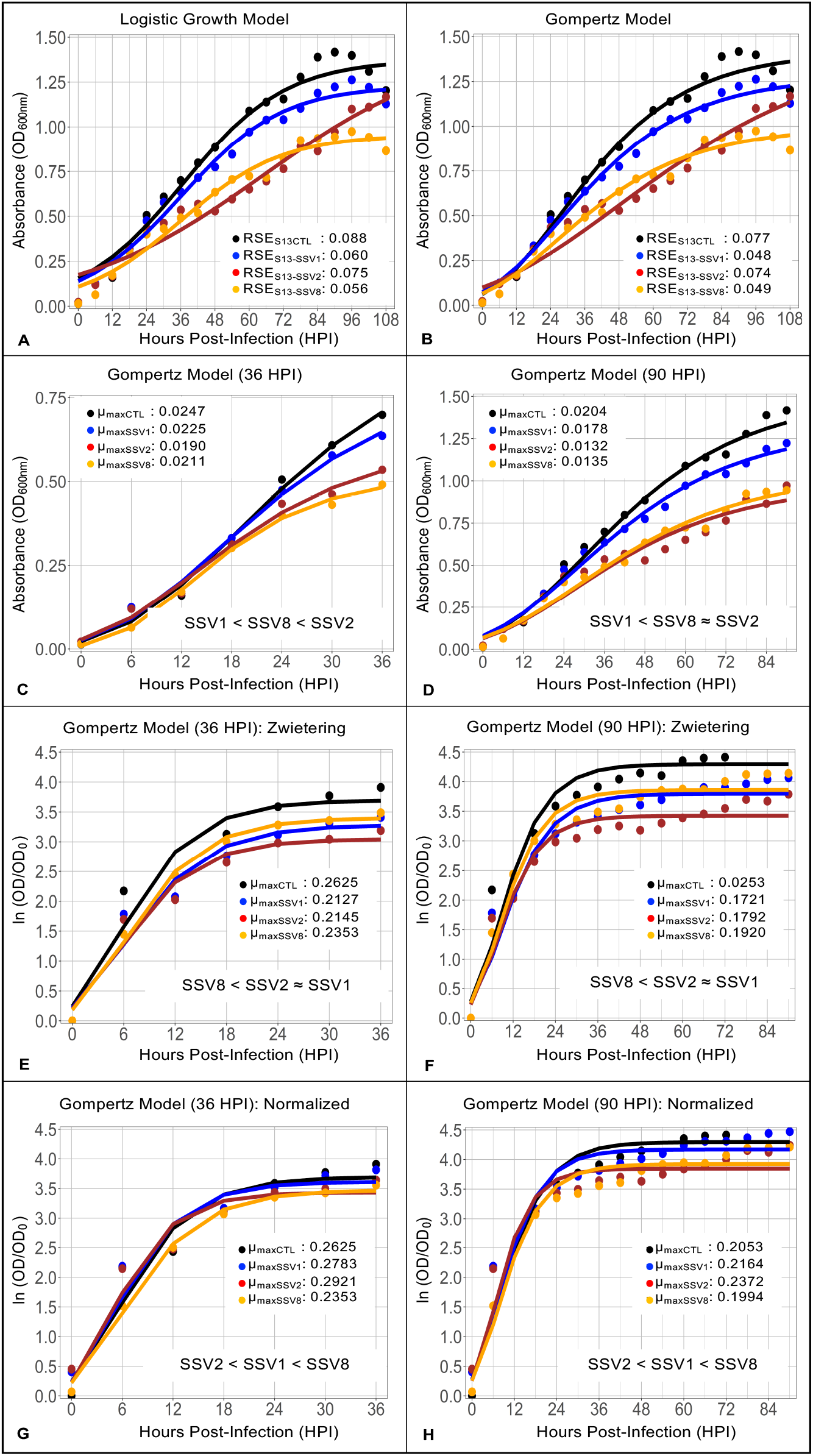
Growth Curve Analysis for SSV Data Using Maximum Specific Growth Rate (μ_max_). Growth curves were generated using host *Sulfolobus* strain Gθ [6,7] infected with SSV1 [3], SSV2 [4], and SSV8 [5], in single-host/single-virus trials at a MOI = 0.1 (78°C, pH 3.2). Growth of uninfected host control (black), SSV1-infected host Gθ growth (blue), SSV2-infected Gθ growth (red), and SSV8-infected Gθ growth (orange). (A) Logistic Growth model fit to the raw growth data with root mean square errors also called residual standard errors (RSE). Gompertz model fit to the raw host Gθ growth data with RSEs. Maximum specific growth rate (μ_max_) of the Gompertz over a narrower range of 0 to 36 HPI representing classical *log phase*. Expanded growth phase to stationary phase (0 to 90 HPI) with μ_max_ values of the Gompertz. Gompertz of log transformed data [12] with μ_max_ values for the truncated growth curve data (0 to 36 HPI). (F) Gompertz of log transformed and normalized data with μ_max_ values for the expanded host growth curve data (0 to 90 HPI). (G) Gompertz of log transformed data with normalization for start point cell density (0 to 36 HPI). (H) Gompertz of log transformed data with normalization for start point cell density (0 to 90 HPI). Based on μ_max_ values for the Gompertz non-log transformed and log transformed model fits, the order of relative virulence for the viruses (SSV1, SSV2, SSV8) under comparison is provided with least virulent to the left and most virulent to the right. There is no agreement between the analytical treatments regardless if truncated or expanded data are used from the growth curves.

## Results and Discussion

### Comparison of host growth using maximum growth rate (μ_max_) as a metric for relative virulence

Two different intervals were considered. First, an interval from 0 to 36 hours post-infection (HPI), which best represents the archetypal “exponential growth phase” [11] was considered (Fig. 1C). Using *μ*_*max*_ from the Gompertz, SSV1 is less virulent than SSV8 while SSV2 is the most virulent (*see* Fig. 1C). Given that host growth subject to non-lytic viral infection does not always exhibit a classical Monodian profile, an outer bound at 90 HPI was used to capture more of the growth curve (Fig. 1D). Calculating *μ*_*max*_ from the Gompertz for this larger portion of the data changes the results. Specifically, SSV8 appears to exhibit approximately equal virulence to SSV2, while SSV1 remains the least virulent (*see* Fig. 1D).

Thus, for non-lytic infections, a significant change in *μ*_*max*_, which drives interpretation of results, can emerge depending on how much of the curve is considered. Depending on culture size and specific virus-host pairing, the truly *exponential* growth phase may be brief with the majority of positive growth comprising the classically described *deceleration*, before *stationary* phase.

A widely used and agreed upon alternative is to calculate *μ*_*max*_ from a log transformed dataset [12]. Calculating *μ*_*max*_ from log transformed data (i.e., ln OD/OD_0_) using narrow (0-36 HPI) and expanded (0-90 HPI) intervals yields another outcome. Using this method, SSV8 appears to be least virulent while SSV2 and SSV1 are roughly equivalent but greater than SSV8 (Fig. 1E, F). Adding an additional normalization step to compensate for slightly different host cell densities measurements at time of viral inoculation (t_0_), yields yet another result for the data (Fig. 1G, H). Notably, *μ*_*max*_ calculated from the Gompertz fit to the normalized log transformed data suggest that SSV2 is least virulent followed by SSV1 with SSV8 exhibiting highest virulence. Remarkably, none of these analytical adjustments for *μ*_*max*_ as the principal parameter for relative virulence captures the known relationship of SSV1, SSV2, and SSV8 virulence on *Sulfolobus* strain Gθ [7,10]. Thus, methods of determining *V*_*R*_, using *μ*_*max*_ as the key parameter, are inadequate.

### Comparison of host growth using area-under-the-curve (AUC) as a metric for relative virulence

Given limitations of *μ*_*max*_ in determining *V*_*R*_ in non-lytic viral infections, an alternative approach is to determine a percent inhibition (PI) based on area-under-the-curve (AUC) [13–16]. Specifically, determining AUC for infected host growth (AUC_infected_) and uninfected control (AUC_CTL_) provides a calculated PI_AUC_ on non-transformed data such that:

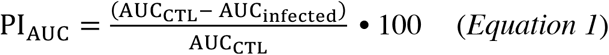

This may be alternatively written as:

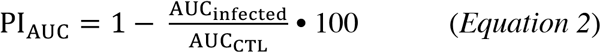

Selecting upper and lower bounds of the integration are critical [17]. Yet, approaches for choosing these bounds are varied and often arbitrary [14,15]. In many reports, the upper bound is chosen absent any noted mathematical or biological explanation. For comparing virus infection impacts, the time of inoculation (t_0_) is a reasonable lower bound; it will capture early changes in host growth. Historically, choice of an upper bound has been subjective. Prior work has relied on a pre-defined *end-point* after culture initiation (e.g., cancer research). A reasonable upper bound is at the beginning of *stationary* phase or *peak growth* (i.e., *N*_*asymptote*_). However, non-canonical host growth during infection may render this value difficult to determine.

Using extremes for the outer limit at 36 HPI and 90 HPI for the *Sulfolobus* strain Gθ-SSV dataset (Fig. 2), AUC is calculated. Given that truly “exponential” growth can be brief for non-lytic infections followed by a long non-exponential growth phase, 36HPI represents a conservative lower limit. Alternatively, the 90 HPI bound captures more of the data extending deeply into the positive growth phase and capturing the growth peak of the uninfected control curve (*see* Fig. 2). Bound at 36 HPI, AUC calculations indicate that SSV1 < SSV2 < SSV8, which is the correct relative virulence between these viruses. However, since this only represents one-third of the dataset, this is only a fortuitous result.

**Fig 2.**
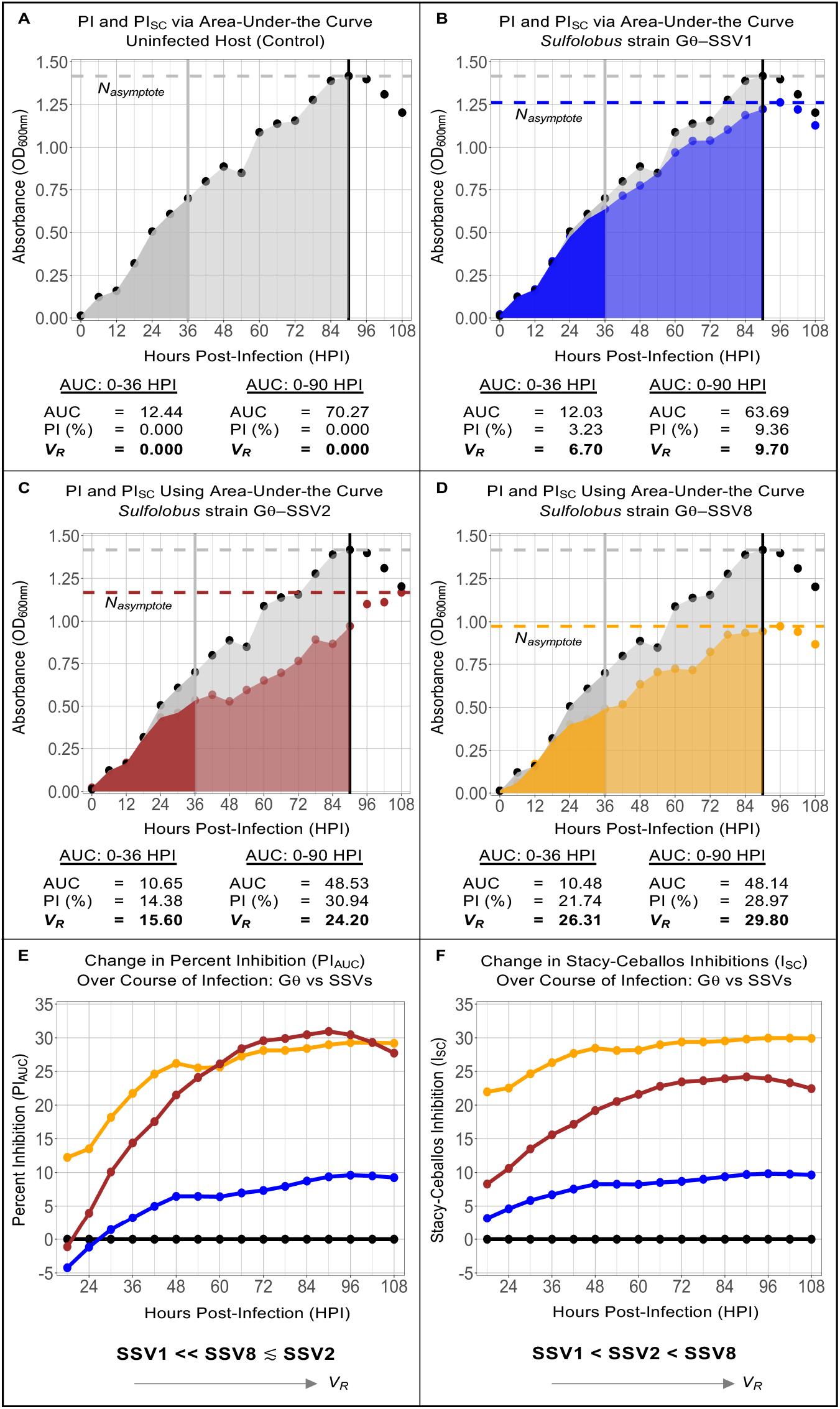
Growth Curve Analysis of SSV Data: Area-Under-the-Curve (AUC). Growth curves for archaeal host *Sulfolobus* strain Gθ [6,7] infected with SSV1 [3], SSV2 [4], and SSV8 [5], in single-host/single-virus trials at a MOI = 0.1 (78°C, pH 3.2). AUC for: (A) uninfected control (black); (B) SSV1-infected host *Sulfolobus* strain Gθ (blue); (C) SSV2-infected *Sulfolobus* strain Gθ (red); and, (D) SSV8-infected host *Sulfolobus* strain Gθ (gold). AUC was calculated for each SSV-infected host growth curve (and uninfected control) using two different sets of integration bounds: 0-36HPI and 0-90HPI. Changes in traditional PIAUC as a function of upper integration bound over the complete dataset with resulting relative virulence (shown at bottom of the panel). Changes in *I*_SC_ as a function of chosen upper bound over the entire dataset with interpretation of relative virulence based on this new approach shown at the bottom of the panel. Regardless of interval used to calculate AUC, using the *I_SC_* approach yields a correct result in terms of relative virulence (*V*_*R*_) while the PI_AUC_ traditional approach does not. All infections were conducted at the same MOI and equivalent starting host cell concentrations since these parameters will influence host growth dynamics during viral infection.

To capture a larger component of the virus-host interaction through the peak growth (*N*_*asymptote*_) of the uninfected control data, the bound was moved to 90 HPI, yielding: SSV1 << SSV8 ≲ SSV2. This is inaccurate and demonstrates that assessing virulence using PI from AUC is unreliable and sensitive to limit selection. What is needed is a metric that captures a significant component of the virus-host interaction (i.e., to peak growth) while yielding the correct *V*_*R*_ between viruses.

### Comparison of host growth using Stacy-Ceballos Equation as a metric for relative virulence

Although *μ*_*max*_ and AUC can be useful parameters for characterizing: drug interactions [18] or attenuated/enhanced growth in mutant versus wild-type cell growth [14,19], when comparing virulence between non-lytic viruses on a host, these parameters are not resilient to changes in integral bounds. An essential component of virus-host interactions is *N*_*asymptote*_ (i.e., peak growth), which is a critical but often ignored parameter in growth curve analysis [17]. By considering both percent inhibition of the growth phase as well as the percent inhibition in *N*_*asymptote*_, a more robust representation of *V*_*R*_ can be determined. Notably, the square root of the product of *PI*_*AUC*_ and *PI*_*max*_, introduced here as *Stacy-Ceballos* inhibition (*I*_*SC*_), provides a robust index of *V*_*R*_, where:

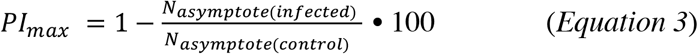

and,

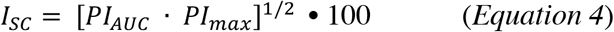

Using *I*_*SC*_, the correct order of increasing virulence emerges for both the 36 HPI and 90 HPI limits (Fig. 2F) with the latter representing a broad range across the virus-host dynamic (*see* Fig. 2A-D). *I*_*SC*_ is a robust index for *V*_*R*_ that is resilient to limit selection. Previous cautions against combining parameters into a single metric are acknowledged [20]; however, combining PI_AUC_ and PI_max_ allows for inclusion of variation between growth curves beyond exponential phase. Furthermore, the use of relative measures allows for meaningful comparisons across virus-host systems. Although growth curves with similar growth patterns will result in an *I*_*SC*_ value similar to PI_AUC_, the former more reliably quantifies differences between growth curves exhibiting distinct growth patterns.

## Conclusions

### Generalizability of Stacy-Ceballos Inhibition (I_SC_) as metric for relative virulence

The non-lytic Sulfolobus Spindle-shaped Virus (SSV) system, provides one example of how traditional parameters for assessing relative virulence (i.e., *μ*_*max*_ and AUC) between two (or more) viruses on a given host may yield unreliable results and incorrect interpretations of infection data. Using Stacy-Ceballos Inhibition (*I*_*SC*_) as a metric for calculating relative virulence overcomes the sensitivity of these parameters providing a more robust and reliable approach for determining *V*_*R*_. Importantly, the robustness and reliability of this approach is not constrained to non-lytic viruses. It is also useful when comparing non-lytic versus lytic infections. In this case, PI_max_ for the lytic system would be the maximum cell density reached prior to lysis. The ability to accurately assess differences in virulence between lytic viruses and non-lytic viruses or changes in virulence as a virus switches between non-lytic (but productive) and lytic phases offers new opportunities in characterizing single-virus/single-host interactions. In a separate report, reliability and robustness of this approach is demonstrated across multiple areas of application [17]. We are also assessing the applicability of *I*_*SC*_ to polymicrobial infections.

